# A Hybrid PINN-DE Framework for Data-Driven Parameter Estimation of Tumor-Immune Dynamics in Bladder Cancer

**DOI:** 10.64898/2026.02.17.706276

**Authors:** Antonio Mastroberardino, Adam Glick

## Abstract

Bladder cancer presents significant clinical challenges due to its complex immune microenvironment and highly heterogeneous response to treatments. To create accurate, individualized models of disease progression, we first construct a system of Ordinary Differential Equations (ODEs) that captures tumor-immune interactions. We address the challenge of estimating unknown parameters by performing a rigorous comparative analysis of two heuristic optimization methods: Differential Evolution (DE), a robust global optimization algorithm, and Physics-Informed Neural Networks (PINN), a novel machine learning framework that embeds ODE constraints into its loss function. Our findings provide a critical evaluation of the computational efficiency and accuracy of each method for parameterizing biological ODE systems. This study validates the power of hybrid machine learning approaches in mathematical oncology, yielding a robust computational framework for parameter estimation and providing a necessary algorithmic foundation for future personalized treatment strategies.

**Author summary:** Bladder cancer remains a major global health threat, characterized by highly unpredictable responses to treatment and a high likelihood of recurrence. To better predict how a patient’s disease will progress, researchers use mathematical models that simulate the interactions between cancer cells and the immune system. However, these models are only useful if they can be accurately tuned to a specific patient’s data—a process called parameter estimation.

This task is notoriously difficult because clinical data is often sparse and noisy, making it hard to find the right settings for the model. In this study, we developed a novel computational framework that combines a traditional optimization algorithm (Differential Evolution) with Physics-Informed Neural Networks (PINNs), a specialized architecture designed to embed physical constraints directly into the learning process.

By “teaching” the AI the underlying biological laws of cancer growth, our hybrid approach can accurately estimate a patient’s unique disease parameters even when raw data is limited. We validated this method using a “virtual patient” system derived from real-world clinical trials. Our results show that this hybrid approach provides a more robust and reliable way to personalize cancer models, offering a powerful new tool for doctors to simulate and optimize individual treatment plans before they are even administered.

## Introduction

Bladder cancer is a significant global health challenge, which ranks as the ninth most common cancer worldwide with approximately 614,000 new cases and 221,000 deaths annually [1]. Incidence and mortality are projected to rise substantially over the coming decades due to aging populations and persistent risk factors such as tobacco exposure. Projections by the World Health Organization indicate that new cases will more than double to over 1.3 million by 2050, with deaths nearly doubling to 440,000 [1]. Clinical management depends on disease stage, classified primarily as non-muscle-invasive bladder cancer (NMIBC), which accounts for approximately 75% of cases, muscle-invasive bladder cancer (MIBC), or metastatic disease [2].

For patients with NMIBC, the primary first-line treatment is transurethral resection of bladder tumor (TURBT) followed by risk-stratified adjuvant intravesical therapy [3]. Low-risk NMIBC is typically managed with a single, immediate intravesical instillation of chemotherapy with either gemcitabine or mitomycin [4–8]. For intermediate- and high-risk disease, the standard first-line treatment is a six-week induction course of intravesical Bacillus Calmette-Guérin (BCG) therapy [9–11].

MIBC and metastatic disease require aggressive systemic interventions, including cisplatin-based chemotherapy, radical cystectomy, or increasingly, immune checkpoint inhibitors (ICIs) such as pembrolizumab and atezolizumab [12–22]. While platinum-based chemotherapy has traditionally been the first-line palliative treatment for metastatic disease, the landscape is rapidly evolving.

Across all disease stages, treatment outcomes vary substantially from patient to patient. NMIBC exhibits recurrence rates as high as 70%, high-risk NMIBC can progress to MIBC, and metastatic disease shows wide variability in response to systemic therapies. This heterogeneity reflects complex dynamics between tumor cells, immune effector populations, and treatment-induced modulation of the microenvironment. Mechanistic mathematical models, particularly systems of ordinary differential equations (ODEs), offer a principled framework to capture these interactions and inform treatment design.

In bladder cancer, ODE models have been applied to study Bacillus Calmette–Guérin (BCG) immunotherapy, elucidating the role of immune effector activation in tumor control and informing optimal dosing schedules [23–28]. Although these models offer valuable insight on the efficacy of treatment and the optimization of dosing schedules, their predictive utility in clinical settings is often limited by uncertainty in key parameters, such as tumor growth rates, immune killing efficacy, and therapy response coefficients.

Reliable parameter estimation in clinical oncology is fundamentally challenged by the nature of available data. Measurements are typically sparse, noisy, and only partially observed. Although tumor burden may be monitored intermittently by imaging, key components of the tumor microenvironment—-such as localized immune cell populations-—are rarely measured longitudinally. These limitations lead to parameter estimation problems characterized by highly non-convex loss landscapes.

Traditional gradient-based optimization methods (e.g., Adam or L-BFGS) often struggle in this setting, particularly due to the stiffness of the underlying biological ODEs, and are prone to convergence to suboptimal local minima, limiting their reliability for clinical inference.

These challenges have motivated the development of alternative approaches for inverse problems in dynamical systems. Recently, Physics-Informed Neural Networks (PINNs) [29–32] have emerged as a powerful framework by embedding ODE constraints directly into the training process. However, standard PINNs rely on gradient-based optimization and may fail to recover accurate tumor–immune dynamics when critical variables—such as immune recruitment rates—are not observed. In contrast, global optimization methods such as Differential Evolution (DE) [33–36] offer robustness against local minima but are often computationally expensive and less effective at refining continuous dynamical trajectories.

In this study, we develop an ODE-based model of tumor–immune interactions in bladder cancer and address the scarcity of clinical data by introducing a Monte Carlo–based data synthesis pipeline that generates realistic longitudinal trajectories conditioned on patient outcomes. Our model includes pharmacokinetic behavior for chemotherapy and immunotherapy using a dynamic function that more accurately represents the absorption and clearance of the drug. To overcome the challenge of parameter estimation, we present a hybrid computational approach that combines the global search capabilities of DE with the physics-constrained, gradient-based learning of PINNs. We establish a robust computational pipeline capable of solving the stiff, nonlinear inverse problems inherent to parameter estimation and perform a comparative analysis by evaluating the accuracy, robustness, and computational performance of DE and PINNs.

This study provides insight into the strengths and limitations of heuristic optimization versus hybrid machine-learning approaches in mathematical oncology. Ultimately, while the data utilized herein is synthetic, the computational advantages demonstrated by the hybrid PINN-DE framework provide a necessary algorithmic foundation. By successfully navigating the mathematical limitations of noise and partial observability, this work bridges a critical gap between theoretical nonlinear dynamics and the future development of data-driven, individualized treatment models in clinical oncology.

## Model Formulation

We develop a deterministic mathematical framework to simulate the interactions between the bladder tumor burden and the host immune response. This model allows for the investigation of tumor progression under natural immune surveillance and the subsequent response to therapeutic interventions.

### Tumor-Immune Dynamics Without Therapy

Let *T*(*t*) and *I*(*t*) denote the tumor cell population and the immune effector cell population (e.g., cytotoxic T-lymphocytes and Natural Killer cells) at time *t*, respectively. We propose a coupled system of ordinary differential equations (ODEs) based on the following biological assumptions:

1. **Tumor Growth:** In the absence of an immune response, tumor cells exhibit logistic growth, characterized by an intrinsic growth rate *r* and a carrying capacity 1*/b*.
2. **Immune Influx and Homeostasis:** The immune population is maintained by a constant source rate *σ* (representing influx from bone marrow or lymph nodes) and decays at a natural death rate *δ*.
3. **Immunosurveillance:** Immune cells eliminate tumor cells upon contact. We model this interaction using Holling Type I functional response with a rate constant *α*_*ti*_.
4. **Immune Recruitment and Inhibition:** At low tumor burden, tumor-associated antigens stimulate immune recruitment at a rate *α*_*it*_. However, increasing tumor burden promotes an immunosuppressive microenvironment. We model this effect phenomenologically by introducing a threshold 1*/β* at which immune recruitment is fully suppressed by the tumor.

The inhibition of the immune system by a tumor is commonly described as immune tolerance [37], during which the immune response becomes incapable of recognizing tumor antigens as foreign or mounting an effective attack. Tumors develop immune tolerance through multiple, overlapping mechanisms that collectively suppress or evade antitumor immunity [38–40]. A common mechanism involves altered antigen expression and presentation, whereby malignant cells downregulate tumor-associated antigens or MHC class I molecules to avoid recognition by cytotoxic T lymphocytes. In parallel, many tumors activate immune checkpoint pathways by overexpressing ligands such as PD-L1 and PD-L2, which bind to PD-1 on effector T cells and transmit inhibitory signals that attenuate proliferation and cytotoxic function.

The tumor microenvironment further reinforces tolerance through immunosuppressive signaling [41, 42]. Tumor and stromal cells secrete inhibitory cytokines such as transforming growth factor-*β* (TGF-*β*) and interleukin-10 (IL-10) to inhibit antigen presentation and effector T-cell activation. These signals also promote the recruitment and expansion of regulatory immune populations including regulatory T cells (Tregs), myeloid-derived suppressor cells (MDSCs), and M2-polarized macrophages, which collectively maintain an anti-inflammatory environment.

Exhausted T cells remain present but express high levels of inhibitory receptors, preventing them from mounting effective responses. Together, these processes work synergistically to create a tolerant immune landscape in which tumors can grow and evolve unchecked.

Fig. 1 presents a schematic illustration of the key biological components included in the model and their interactions. The governing system of ODEs based on these assumptions is given by

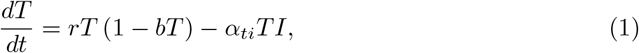

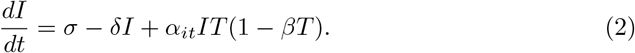

**Fig 1.**
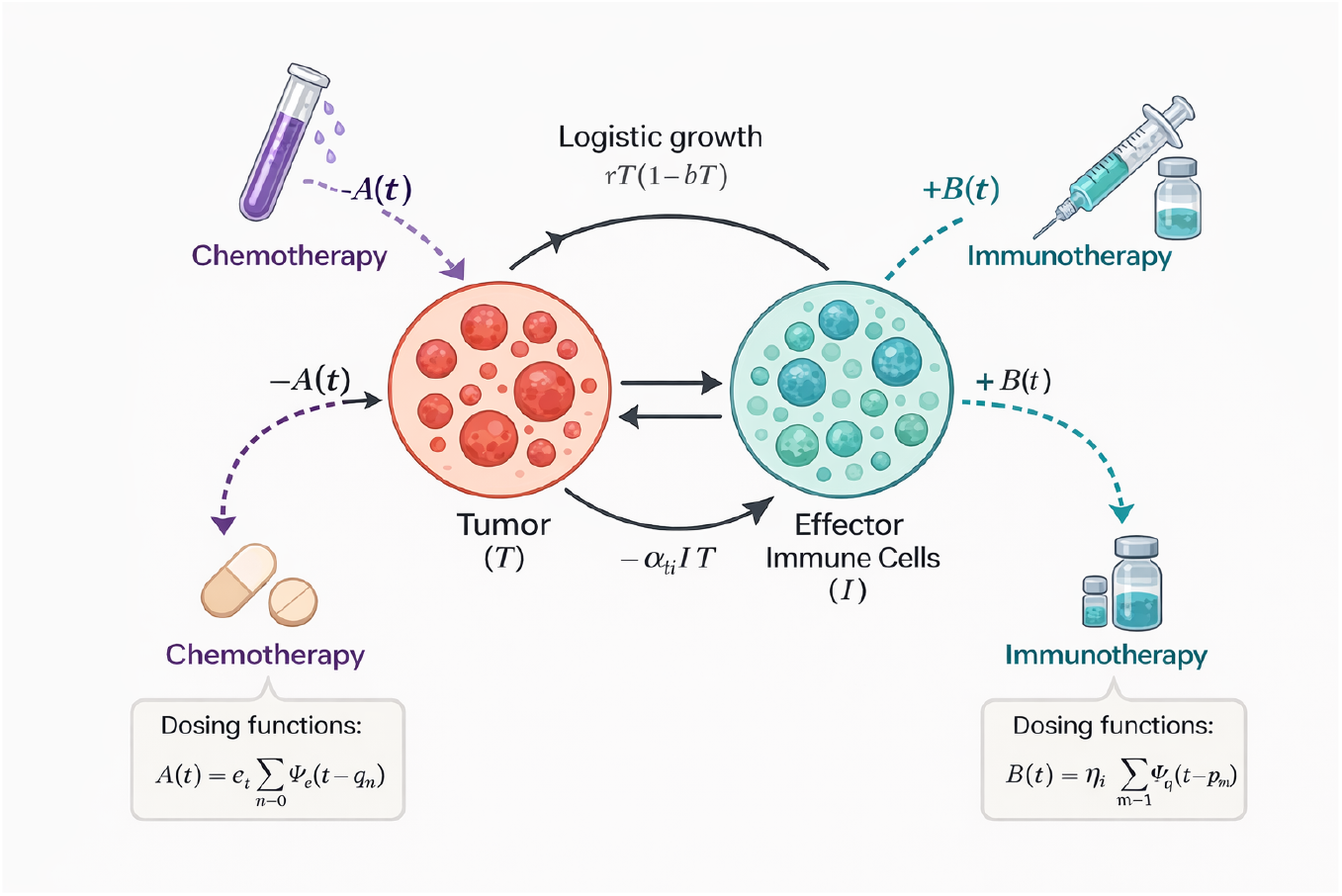
Schematic representation of the tumor–immune interaction model with therapeutic inputs. Tumor cells undergo logistic growth and interact with immune cells through cytotoxic and stimulatory mechanisms. Chemotherapy induces tumor cell death, while immunotherapy enhances immune response to restore antitumor activity and reduce tumor burden. Time-dependent treatment effects are modeled through convolution-based dosing functions. Image created at bioRender.com.

See Table 1 for a full description of the system parameters.

**Table 1.** Model parameters and biological interpretation.

| Parameter | Units | Description |
| --- | --- | --- |
| $r$ | $\text{week}^{-1}$ | Intrinsic tumor growth rate |
| $b$ | $\text{cells}^{-1}$ | Tumor carrying capacity parameter ($1/b$) |
| $\alpha_{ti}$ | $\text{cells}^{-1} \text{week}^{-1}$ | Immune-mediated tumor killing rate |
| $\sigma$ | $\text{cells week}^{-1}$ | Constant influx rate of immune effector cells |
| $\delta$ | $\text{week}^{-1}$ | Natural death rate of immune effector cells |
| $\alpha_{it}$ | $\text{cells}^{-1} \text{week}^{-1}$ | Tumor-induced immune recruitment rate |
| $\beta$ | $\text{cells}^{-1}$ | Immune inhibition (tolerance) strength |
| $\epsilon$ | $\text{week}^{-1}$ | Maximal chemotherapy-induced tumor kill rate |
| $\eta$ | $\text{week}^{-1}$ | Maximal immunotherapy-induced immune activation rate |
| $\omega_\epsilon$ | week | Chemotherapy absorption half-life |
| $\tau_\epsilon$ | week | Chemotherapy clearance half-life |
| $\omega_\eta$ | week | Immunotherapy absorption half-life |
| $\tau_\eta$ | week | Immunotherapy clearance half-life |
| $C_\epsilon$ | – | Chemotherapy dose normalization constant |
| $C_\eta$ | – | Immunotherapy dose normalization constant |

### Model with Therapeutic Intervention

We now include therapy terms in equations (1)−(2) to model a realistic administration of immunotherapy and chemotherapy. To account for the fact that clinical dosing is discrete and transient, we model the *effective drug concentration* at the tumor site using pharmacokinetic decay functions.

Let *A*(*t*) and *B*(*t*) represent the time-dependent efficacy of chemotherapy and immunotherapy, respectively. This formulation is mathematically equivalent to the convolution-based impulse–response model introduced in [43], but is written here in a form that explicitly highlights treatment accumulation and clearance. The extended system is defined as

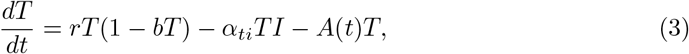

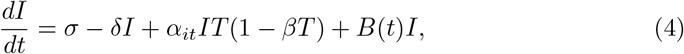

where the chemotherapy term −*A*(*t*)*T* represents the direct fractional kill of tumor cells, while the immunotherapy term +*B*(*t*)*I* represents the enhanced proliferation or reduced exhaustion of effector cells.

Following the approach proposed by Glick and Mastroberardino [43], we model the drug action not as a constant, but as a dynamic function of uptake and clearance, thereby providing a more accurate representation of the drug’s pharmacokinetics. For a single dose administered at time *t* = 0, we define the efficacy functions for chemotherapy and immunotherapy, respectively, as

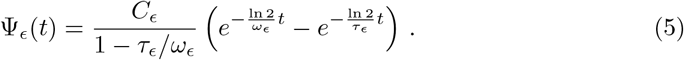

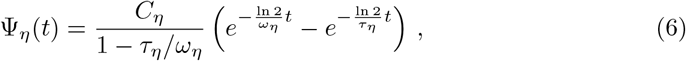

where *ω*_*i*_ is the effective absorption half-life, *τ*_*i*_ is the decay half-life, and *C*_*i*_ is a normalization constant regulating dose intensity for *i* = *ϵ, η*.

For a treatment schedule consisting of *N* chemotherapy doses administered at times {*q*_0_, *q*_1_, …, *q*_N−1_} and *M* immunotherapy doses at times {*p*_0_, *p*_1_, …, *p*_M−1_}, the total therapeutic functions *A*(*t*) and *B*(*t*) are constructed via the summation of shifted efficacy functions:

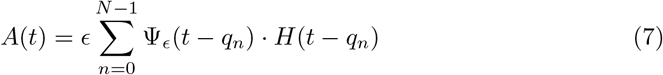

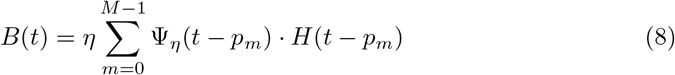

where *H*(·) is the Heaviside step function, and *ϵ* and *η* scale the maximal potency of chemotherapy and immunotherapy, respectively. This formulation ensures that biological outcomes depend on the realistic accumulation and clearance of therapeutic agents while remaining compatible with gradient-based and physics-informed parameter estimation techniques.

### Computational Formulation

Robust training of PINNs requires an adequate amount of data in order to obtain meaningful results. However, public datasets for bladder cancer typically report aggregate survival statistics or discrete response rates per the RECIST criteria [44] rather than raw, time-series cellular measurements. To overcome this data scarcity, we develop a Monte Carlo simulation framework to synthesize longitudinal patient trajectories based on clinical trial meta-data reported by Song et al. [28].

We synthesize data using a custom built Monte Carlo script that randomly creates cell population data in the form of a time series sampled within the bounds of clinical trial results. The synthetic data is then used as the basis for parameter estimation using DE and PINNs to fit the data to Eqs. (3) – (4). All software used to generate the results presented in this work, including the code used to simulate the data, is publicly available at [45].

### Data Synthesis Methodology

We synthesize data by aggregating clinical trials presented in Song *et al*. [46] and summarized in Table 2. The published dataset contained only aggregate data and no individual patient information. Leveraging this data, we develop a Monte Carlo simulation in Python to generate longitudinal tumor cell count data for bladder cancer patients across various clinical trials. Each trial was defined by its unique identifier, drug, drug category, and empirical response rates, described as Complete Response (CR), Partial Response (PR), Stable Disease (SD), or Progressive Disease (PD). A total of 13 trial arms were simulated and patients were probabilistically assigned one of the four response outcomes using a multinomial distribution based on observed clinical response rates. For reproducibility, a fixed random seed was applied to a random number generator.

**Table 2.** Clinical trials aggregated by Song *et al*. [46], showing the clinical trial number, number of enrolled patients, therapy modality, and drug used in each study. The chemotherapy drugs used in all cases were paclitaxel, docetaxel, or vinflunine. Only aggregate data were reported; no individual patient data were included in the original publication.

| Clinical Trial | # of Patients | Type of Therapy | Drug Name |
| --- | --- | --- | --- |
| NCT01375842 | 68 | Immunotherapy | Atezolizumab |
| NCT02108652 | 310 | Immunotherapy | Atezolizumab |
| NCT02951767 | 119 | Immunotherapy | Atezolizumab |
| NCT02302807 | 466 | Immunotherapy | Atezolizumab |
| NCT02302807 | 466 | Chemotherapy | Paclitaxel, Docetaxel, or Vinflunine |
| NCT01772004 | 293 | Immunotherapy | Avelumab |
| NCT01928394 | 78 | Immunotherapy | Nivolumab |
| NCT02387996 | 265 | Immunotherapy | Nivolumab |
| NCT01848834 | 33 | Immunotherapy | Pembrolizumab |
| NCT02256436 | 271 | Immunotherapy | Pembrolizumab |
| NCT02256436 | 271 | Chemotherapy | Paclitaxel, Docetaxel, or Vinflunine |
| NCT02335424 | 370 | Immunotherapy | Pembrolizumab |
| NCT02736266 | 50 | Immunotherapy | Pembrolizumab |

Tumor burden is modeled as a proxy for cell count, derived from tumor diameter. We assume tumor mass scales with volume and spherical geometry of the tumor so that the cell count *N* (*t*) is proportional to the cube of the tumor diameter *D*(*t*). The RECIST threshold for partial response of 30% reduction in diameter corresponds to a volumetric reduction of:

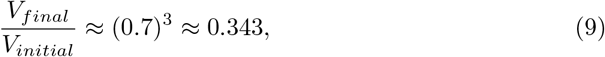

which implies a reduction in tumor cell count of approximately 65.7%.

We model the temporal dynamics of the tumor count for responders using a stochastic exponential decay model. For CR patients, the tumor cell count exhibits exponential decay that approaches zero defined by:

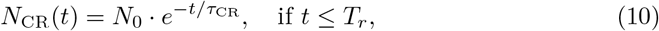

where *N*_0_ = 10^6^ is the initial tumor cell count, *T*_*r*_ *≈* 6 weeks is the response time, and

*τ*_CR_ is a personalized decay constant such that

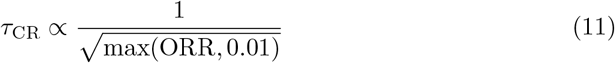

where ORR is the trial’s objective response rate. For PR patients, cell counts also decay exponentially but stabilize at a nonzero target, sampled from a normal distribution centered at *µ* = 3.43 *×* 10^5^ with standard deviation *σ* = 0.1*µ*:

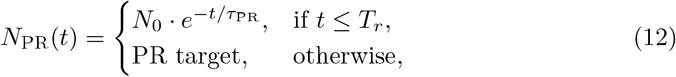

where

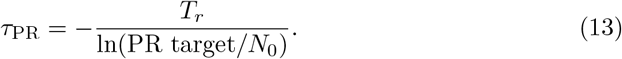

The Python script is designed to support Monte Carlo generation of multiple synthetic datasets by seeding the NumPy random number generator with a configurable integer seed at the beginning of each simulation. This ensures both statistical variation and reproducibility. A single call to the script produces one or multiple realizations of the dataset depending on the parameters passed to it. Each seed samples new patient response types, target cell counts for PR, and randomized decay constants. This robust dataset was then used as the basis for data-driven statistical and machine learning techniques to determine the parameters in Eqs. (3) – (8) and test their results.

### Assumptions and Clinical Data Limitations

While the model employs a stochastic framework for data synthesis, the following assumptions are necessitated by the nature of current clinical reporting standards:

- Geometric Linear-to-Volumetric Mapping: We acknowledge that real-world tumors rarely exhibit perfect spherical symmetry. However, in the absence of routine 3D volumetric imaging in large-scale clinical trials, we adopt the standard RECIST 1.1 assumption. By modeling tumor burden as proportional to the cube of the longest diameter (*D*^3^), we maintain consistency with how clinical “Partial Response” and “Progressive Disease” are defined globally. This approximation serves as a necessary bridge between 1D clinical measurements and 3D biological dynamics.
- Data-Driven Proxy for Cell Count: True tumor cell counts (*N*) are not observable in living patients. We utilize the available longitudinal diameter data as a primary proxy, assuming a constant cellular density. While this omits complexities like necrotic centers or stromal infiltration, it provides a mathematically tractable baseline that honors the signal-to-noise ratio found in actual patient records.
- Hybrid Decay-Growth Dynamics: Critics may note the use of exponential decay in the synthesis phase; however, this is a localized approximation of the initial treatment response phase observed in the clinical data. To account for long-term biological realism, these trajectories are integrated with logistic growth constraints (smeared via randomized parameters). This ensures that the synthetic data reflects the biological reality of carrying capacity and competitive inhibition, rather than unbounded growth or decay.
- Source Data Constraints: The synthetic dataset is grounded in the limited pool of available longitudinal human trials. The decision to use a stochastic Monte Carlo approach was made specifically to address the “sparsity problem” in oncology data. By seeding the generator with randomized decay constants and target counts, we capture the inter-patient variability that a purely deterministic, perfectly “spherical” model would miss.
- Focus on Framework over Morphological Fidelity: The primary goal of this research is to validate the parameter estimation framework. While non-spherical volumes and non-exponential kinetics exist, the methodology demonstrated here is designed to be “physics-informed” and agnostic to the specific growth law; it can be recalibrated as higher-fidelity volumetric data becomes available in the future.

### Parameter Estimation via Differential Evolution

We employ DE to establish a baseline for parameter estimation and provide robust initialization for the PINN. DE is a stochastic, population-based global optimization algorithm that excels in non-convex, non-differentiable landscapes common in biological inverse problems. Unlike gradient-based methods, which are prone to entrapment in local minima, DE evolves a population of candidate parameter vectors through mutation, crossover, and selection. This allows for an extensive exploration of the parameter space before refining the solution.

We identify a set of parameters ***θ*** that minimize a composite loss function ℒ(***θ***), which quantifies the discrepancy between the ODE model predictions and the observed synthetic data. Optimization is carried out in an unconstrained parameter space ***ϕ***, where physical parameters are recovered via exponential mapping and constrained within biologically plausible ranges. Therapy-specific scenarios are handled by setting degenerate bounds (e.g., *ϵ*_*t*_ = *τ*_*ϵ*_ = *C*_*ϵ*_ = 0 in immunotherapy-only cases), allowing a single objective formulation to flexibly adapt across treatment regimes.

### DE Implementation and Objective Formulation

The DE optimizer maintains a population of **M** candidate solutions, each of dimension **D**, where **D** is the number of parameters to be estimated. The population matrix, **P**, can be represented as a stack of row vectors 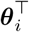, defined as

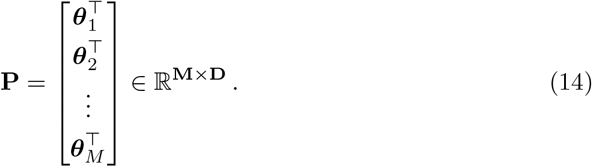

Mutation in differential evolution generates a mutant vector for each candidate as

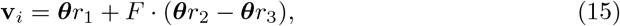

where *r*_1_, *r*_2_, *r*_3_ ∈ 1, …, **M** are distinct indices and *F >* 0 is the mutation factor controlling the step size.

Crossover produces a trial vector **u**_*i*_ by mixing components of the mutant vector with those of the target vector. For each component *j* ∈ 1, …, **D**, draw a random number

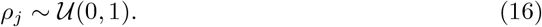

The trial vector is then defined as

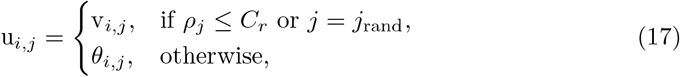

where *C*_*r*_ ∈ [0, 1] is the crossover probability and *j*_rand_ is a randomly chosen index ensuring that at least one parameter component is inherited from the mutant vector. This balance of mutation and crossover enhances both global exploration and local refinement in the parameter estimates.

Evaluations of the objective function are parallelized using Python’s **multiprocessing** module, allowing multiple candidate solutions to be evaluated simultaneously. A shared-memory array tracks the current best solution, and a lock ensures thread-safe updates, which triggers the callback function for early stopping when appropriate. Callback functions allow early termination once a target loss or computational budget is reached, and a fixed random seed ensures reproducibility of the differential evolution runs.

For a given candidate ***θ***, the scaled tumor–immune system is integrated numerically using **scipy.integrate.solve_ivp** with the LSODA method, which adaptively switches between stiff and non-stiff solvers. To prevent differential evolution from diverging due to excessively large tumor and immune populations as well as preventing overflow errors in our computations, the tumor and immune populations are normalized by scale factors

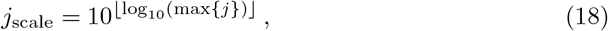

where *j* ∈ {*T, I*} so that

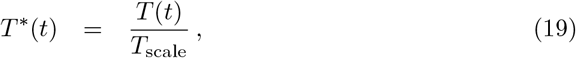

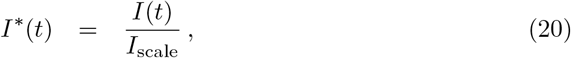

with initial conditions set from the first observed tumor count and mean immune count. After integration, predictions are rescaled to get the model’s full result as

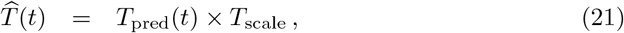

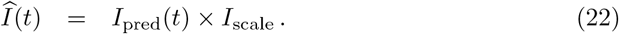

To account for inter-patient variability, the loss is computed as an average across all patients (**N**) and evaluation times (**M**) taking inspiration from [47], [48], and [49]:

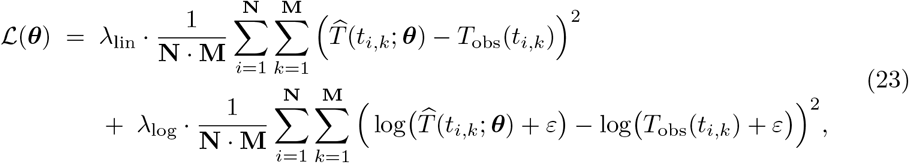

Here, the linear-space term emphasizes absolute deviations at larger tumor burdens, while the log-space term emphasizes relative deviations during early growth or regression phases. The small constant *ε >* 0 prevents divergence as tumor sizes approach zero. For this work, we do an average of the two loss terms by setting *λ*_lin_ = *λ*_log_ = 0.5 with a constant value *ε* = 10^−2^.

Finally, if the ODE solver fails due to stiffness, instability, or unbounded growth, or if parameters fall outside admissible ranges, the objective function is assigned a large penalty (e.g., 10^6^), thereby steering the optimizer away from infeasible regions of parameter space.

### Parameter Constraints

To enforce positivity and stabilize the optimization, the optimizer operates on unconstrained parameters ***ϕ***, with the physical parameters recovered inside the loss function via

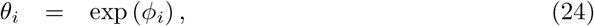

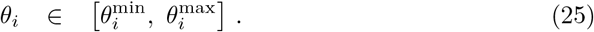

This enforces positivity and constrains the estimated parameters within user-specified bounds.

The optional upper and lower bounds are passed to the DE optimizer via the **bounds** argument for additional safety. These bounds are also enforced inside the Python objective function via clamping, providing a second layer of protection against invalid parameters. The bounds chosen are specific to each parameter and typically spanning several orders of magnitude to allow exploration of wide parameter regions and identification of optimal values.

### Parameter Refinement with PINN

Following coarse parameter estimation via DE, we refine the model using PINN. The DE procedure produces an optimized parameter set ***θ***^*^ that serves as the initialization for PINN. This global search reduces the risk of poor local minima and improves the likelihood of biologically meaningful convergence during PINN training. The PINN simultaneously fits tumor-immune dynamics to the data while enforcing the underlying ODE structure, enabling accurate interpolation and biologically consistent extrapolation.

### Model Architecture and Loss Formulation

The data generated in Sect. is randomly separated into training and testing data using a random permutation of indices across the dataset. First, a seed is set for permutation reproducibility so that the result of the PINN’s model and parameter estimations can be assessed during hyperparameter tuning. The training set is passed into the training loop and the testing set is used to assess the PINN’s performance at estimating data evolution in time. For numerical stability, observed tumor and immune cell populations are normalized by their maximum observed values before training. This prevents gradients from exploding to positive and negative infinity, allowing training to commence without encountering overflow errors.

Let 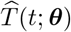 and 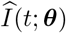 denote the neural estimates of tumor and immune cell populations over time using PINN. The network is parameterized by trainable weights and biases (collectively denoted by ***θ***) as well as system parameters appearing in the governing ODEs. The PINN receives as input a time point *t* and outputs estimates of both *T* and *I*. To enforce the ODE constraints, we require that

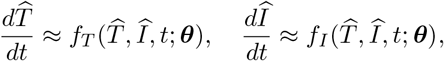

where *f*_*T*_ and *f*_*I*_ are the right-hand sides of Eqs. (3) – (4).

The network mapping is defined as

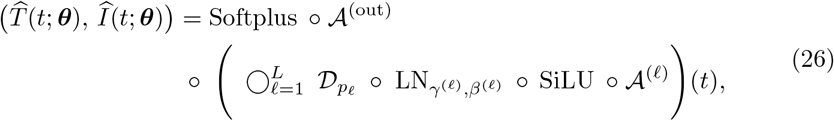

with nodes *n*_0_ = 1, *n*_*ℓ*_ = 200 for *ℓ* = 1, …, *L, n*_L+1_ = 2, *L* = 6 hidden layers, and 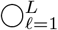 represents function composition in sequence defined by

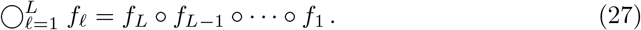

Each hidden layer consists of four operations applied sequentially. First, an *affine transformation* is applied,

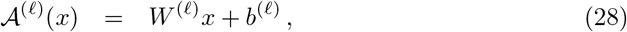

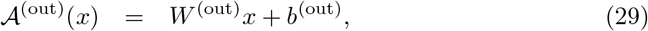

with weight matrices *W* ^(*ℓ*)^ and biases *b*^(*ℓ*)^ as trainable parameters. These are followed by the *SiLU activation* (also known as Swish-1),

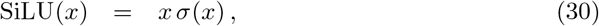

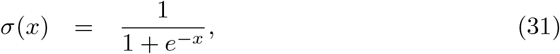

which introduces smooth nonlinearity. To improve stability during training, a *layer normalization* step rescales and shifts the activations,

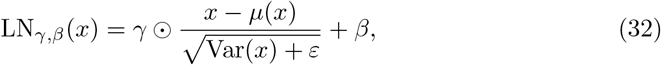

where *γ, β* ∈ ℝ^200^ are trainable parameters, *ε >* 0 is a small constant, and *µ*, Var denote the feature mean and variance. ⊙ is the Hadamard multiplication operator where if *A, B* ∈ ℝ^*m*×*n*^,

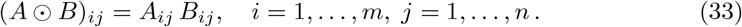

Finally, *dropout* is applied for regularization:

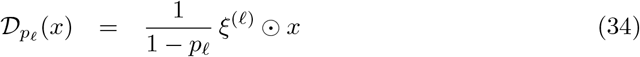

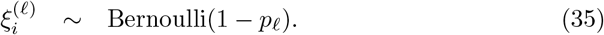

At inference time, dropout reduces to the identity unless Monte Carlo dropout is explicitly enabled.

At the network output, an affine transformation produces a two-dimensional vector *z* = (*z*_1_, *z*_2_), which is then passed through the *Softplus* function to enforce positivity,

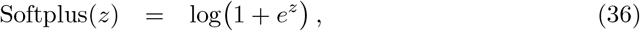

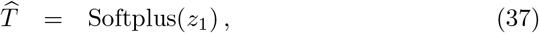

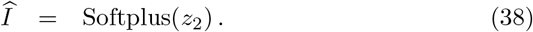

Collecting all trainable weights and biases, the parameter set is

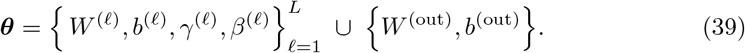

For a vector of times *t* ∈ ℝ^*T*×1^, the mapping acts row-wise, yielding 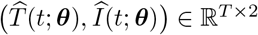.

The total loss function is defined by

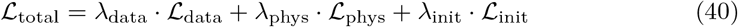

with components

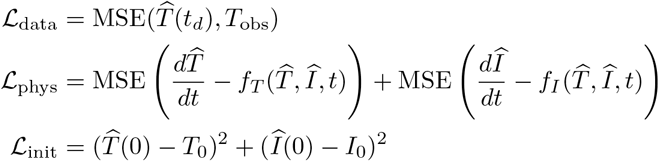

where *t*_*d*_ are a randomly sampled subset of the observed data times. ℒ_data_ is calculated for mini-batches of patients, and *T*_0_, *I*_0_ are the known initial conditions. The physics loss is computed at randomly sampled subset of a fine-grained time discretizations covering *t*_0_, …, *t*_*f*_, often chosen adaptively to focus on high-residual regions.

We start with *λ*_*j*_ values defined in Table 3. The annealing schedule for the physics loss weight, *λ*_phys_, is defined as follows, where *e* is the current epoch, *e*_start_ is the ramp-up start epoch, *e*_end_ is the ramp-up end epoch, *λ*_phys, initial_ is the initial physics loss weight, and *λ*_phys, final_ is the final physics loss weight:

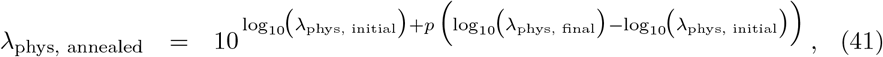

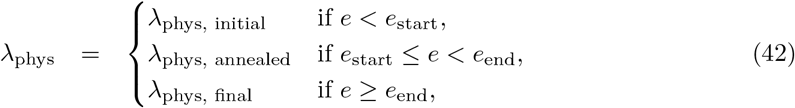

where the progress, *p*, is defined as:

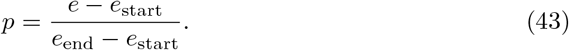

**Table 3.**
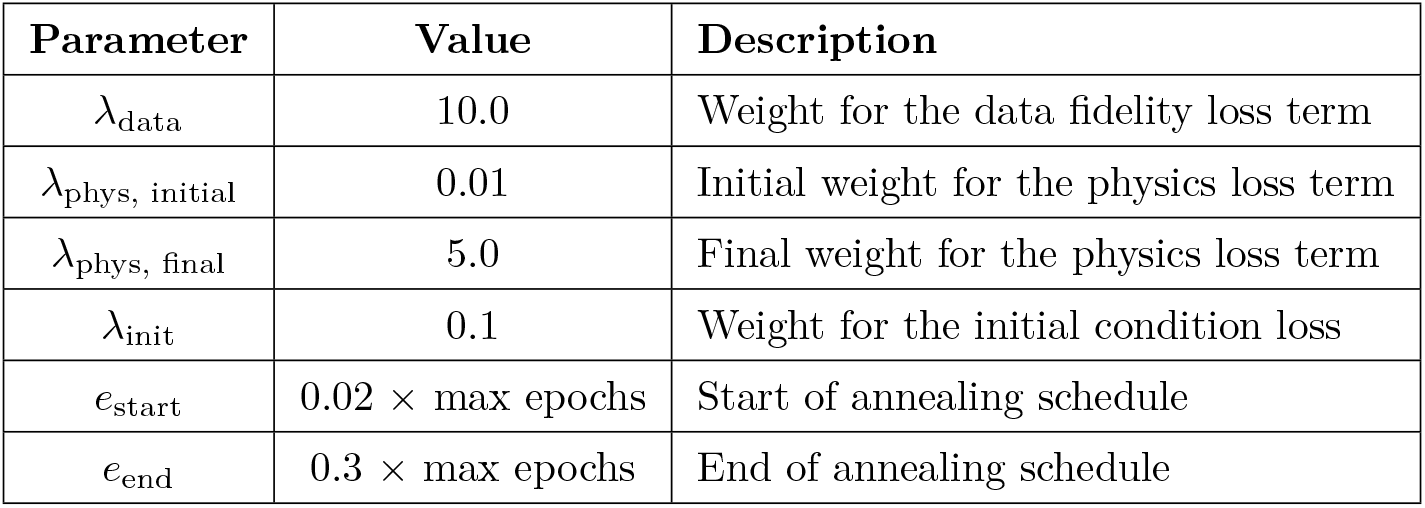
Lambda weight and annealing schedule values for PINN loss terms.

| Parameter | Value | Description |
| --- | --- | --- |
| $\lambda_{\text{data}}$ | 10.0 | Weight for the data fidelity loss term |
| $\lambda_{\text{phys, initial}}$ | 0.01 | Initial weight for the physics loss term |
| $\lambda_{\text{phys, final}}$ | 5.0 | Final weight for the physics loss term |
| $\lambda_{\text{init}}$ | 0.1 | Weight for the initial condition loss |
| $e_{\text{start}}$ | $0.02 \times \text{max epochs}$ | Start of annealing schedule |
| $e_{\text{end}}$ | $0.3 \times \text{max epochs}$ | End of annealing schedule |

Unlike a conventional PINN that treats system parameters as known constants, we treat ***θ*** as trainable variables. Parameters ***θ*** are initialized using the optimized values from the DE procedure, ensuring that training starts in a biologically plausible basin.

To ensure positivity and boundedness of parameters during training, we reparameterize unconstrained network weights through smooth transformations. Common choices include the exponential mapping, which enforces strict positivity, and the logistic sigmoid mapping, which constrains values to a fixed interval. In this work, we adopt the logistic sigmoid because it offers several advantages: it smoothly maps from the entire real line to a bounded interval, preventing runaway parameter growth; its derivatives are well-behaved and do not explode as in the exponential case; and it allows for explicit control of parameter ranges via (*θ*_min,*i*_, *θ*_max,*i*_). This bounded, differentiable mapping improves numerical stability and ensures that gradient-based optimization proceeds more reliably. The sigmoid reparameterization is defined as

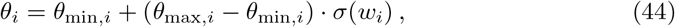

where *w*_*i*_ is the unconstrained network parameter and *σ* is the logistic sigmoid.

### Numerical Implementation and Optimization

Model optimization is performed via automatic differentiation (e.g., PyTorch **autograd**), allowing for the simultaneous update of neural network weights and ODE parameters. Training utilizes the Adam optimizer, employing a bifurcated learning rate strategy where weight decay is applied exclusively to the network weights to prevent overfitting, while ODE parameters are left unregularized to ensure unbiased physical estimation. To maintain a *C*^∞^ differentiable loss landscape and prevent gradient vanishing, discontinuous dose-response functions are replaced with smooth approximations, such as sigmoid-smoothed Heaviside functions. We eschew piecewise linear activations like ReLU—which possess vanishing second derivatives—in favor of smooth transcendental functions. Given the high cardinality of the training dataset and the fine discretization of the temporal vector, we utilize a mini-batch training regime. To ensure global convergence across the entire temporal domain, a stochastic collocation strategy is employed, where a new subset of residual points is resampled every *N* epochs. This approach mitigates the stability issues often associated with mini-batching in PINNs while ensuring that the network adequately captures the transient dynamics of the tumor–immune interactions.

Glick *et al*. [50] proposed a stopping condition for Maximum Likelihood Expectation Maximization (MLEM) that determines the maximum likelihood of the estimation by locating the inflection point of the divergence of the estimation between consecutive iterations. Eq. (47) shows this stopping metric, which can also be used for early termination in PINN training, where 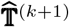 is a vector representing the model’s estimation of the data for all time points on the current epoch with the current parameter estimations, ***θ***^(*k*+1)^, and 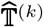 is a vector representing the model’s estimation of the data for all time points on the previous epoch with the previous parameter estimations, ***θ***^(*k*)^.

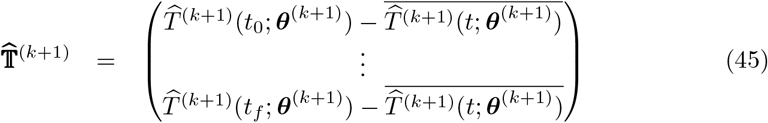

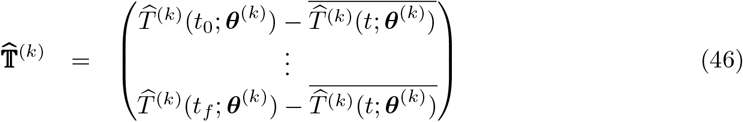

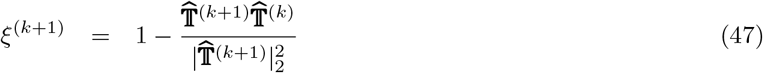

*ξ*^(*k*+1)^ is a akin to a decorrelation measure between consecutive predictions in the tumor count. Unlike in [50] where MLEM was optimized, 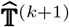 is an estimation from PINN and, therefore, is not a continuously decreasing function with increasing iteration. Eq. (48) is therefore not a valid stopping condition for PINN training in this work. We have constructed a new stopping condition to define the optimal performance of the neural network, which occurs when Eq. (49) is satisfied. To ensure that we do not prematurely break the training loop early, the network training is only halted if both Eq. (49) and Eq. (50) are satisfied for a sufficient number of epochs.

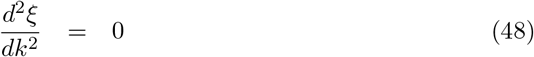

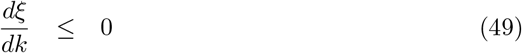

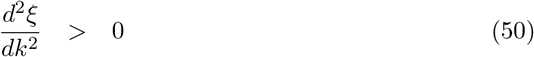

Requiring Eq. (50) to be strictly positive is a necessary condition for early stopping due to the momentum of the decorrelation measure. If the curvature of *ξ*^(*k*+1)^ becomes negative, then the decorrelation is losing momentum and the neural network is heading toward instability between epochs rather than a plateau in its training.

Eqs. (49) – (50) can be combined to create an indicator function that has the form

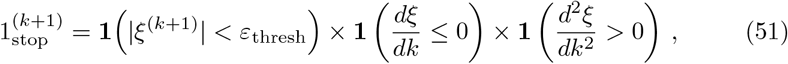

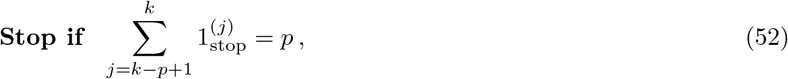

where *ε*_thresh_ is a threshold value to ensure that the decorrelation between estimations is sufficiently small and early stopping is triggered if Eq. (51) holds for *p* consecutive epochs.

The threshold *ε*_thresh_ is determined rigorously by examining how the PINN prediction decorrelated from the DE prediction with increasing epoch using Eq. (47), where 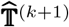 is the tumor count estimated by the PINN at epoch *k* + 1 and 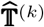 is the optimized DE estimate at epoch *k*. Let 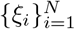 be the decorrelation metrics across all *N* epochs. Then the confidence threshold *ε*_thresh_, corresponding to a confidence level 1 *− α* (we take *α* = 0.01 for 99% confidence), is given by the quantile (inverse cumulative distribution function) of the distribution of |*ξ*|:

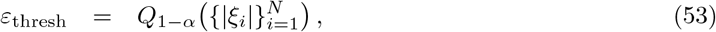

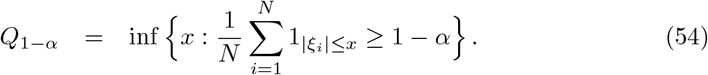

Here, 1_{·}_ denotes the indicator function. For the data used in this work, this procedure yields

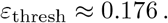

To determine *p*, we assume independence per epoch. We let the event that the stopping condition holds in a single epoch be defined in Eq. (51). If the stochastic fluctuations in training are approximately independent between consecutive epochs, then the probability of a false stop decreases multiplicatively with the number of consecutive epochs required. Therefore the probability of triggered early stopping spuriously over *p* consecutive epochs is

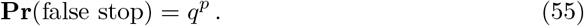

If we want the probability of a spurious stop to be *α*_false_ ≤ 1%, then we get an equation for patience, *p*, that is

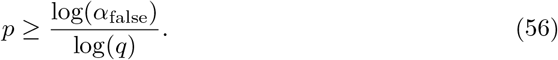

The parameter *q* represents the probability of a “spurious stop” before the network has reached a true plateau. To ensure a robust estimation, we define *q* empirically based on the variance observed in the Differential Evolution (DE) optimization phase:

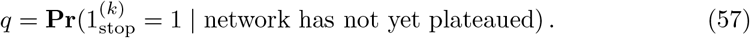

By characterizing the stochastic fluctuations inherent in the DE-derived parameter space, we obtained a value of *q* = 0.240. Substituting this into Eq. (56) yields *p* ≈ 3. This selection of patience ensures that the stopping condition is grounded in the physical variability of the system rather than arbitrary heuristics.

### Uncertainty Quantification via Monte Carlo Dropout

The trained PINN yields both a refined set of parameters ***θ***^⋆^ and continuous, differentiable estimates of the system states over time, 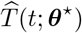 and 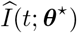. These estimated trajectories provide a smooth functional representation of the underlying dynamics and can be used in an independent numerical integrator (e.g., SciPy’s ODE solver) to verify that the inferred parameters reproduce the observed system behavior, denoted as ***θ***^⋆^ → *T* (*t*) and *I*(*t*). This hybrid approach preserves the interpretability of mechanistic models while leveraging the expressive capacity of neural networks, enabling accurate reconstructions under sparse or noisy observations.

To assess model generalization and estimate predictive uncertainty, Monte Carlo dropout was employed during evaluation. Specifically, dropout layers were explicitly reactivated at test time, and the trained network was sampled multiple times (*S* = 10^3^ forward passes) over a uniform test grid. Each forward pass corresponded to a stochastic realization 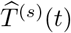 arising from a different random subset of the learned network weights being active. The ensemble of predictions 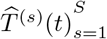 was then used to compute both the mean trajectory 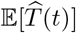 and the associated pointwise variance Var 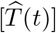, providing an empirical measure of epistemic uncertainty.

This Monte Carlo dropout procedure introduces a Bayesian interpretation to the PINN’s deterministic inference, allowing for statistically valid uncertainty quantification without requiring full variational Bayesian training. The resulting predictive distribution reflects both the robustness of the learned dynamics and the confidence of the model in regions of the state space that were weakly constrained by data.

To quantify the width of the pointwise confidence intervals, we computed the normalized band width (NBW), defined as the ratio of the interval width to the range of the mean trajectory:

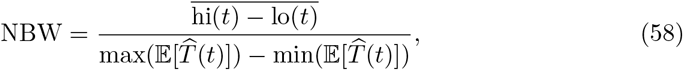

where hi(*t*) and lo(*t*) denote the upper and lower bounds of the 95% pointwise confidence interval at each time point, and the overline indicates averaging over all time points. NBW thus provides a compact, interpretable metric to assess the typical relative width of the Monte Carlo-derived predictive uncertainty.

## Results

Clinical trials taken from [46] were used to synthesize cell data for the complete response and partial response cohorts as previously described. The synthetic cell data was then used to perform parameter evaluation via differential evolution and PINN.

The two methods are compared against the raw data for accuracy of the model’s prediction of the parameters. The two metrics used to determine a model’s predictive accuracy were R^2^, defined in Eq. (59), and mean squared error (MSE), defined in Eq. (60), where *y*_*i*_ is the observed data, 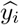 is the model’s prediction of the data, and 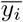 is the mean of the data.

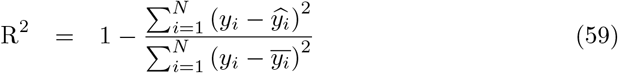

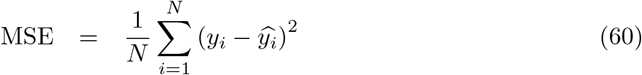

### Complete Response Cohort

Fig. 2 shows the evolution of the conversion metric, *ξ*^(*k*+1)^, plotted alongside a rolling average window taken over approximately 1% of the iterations taken during training and training was ceased after 20,378 epochs, when Eq. (52) was satisfied. *ξ*^(*k*+1)^ shows high divergence between estimations in the early training history, with highly oscillatory behavior in the rolling average window, but as the epochs move forward in time *ξ*^(k+1)^ stabilizes and the rolling average window flattens. Parameter estimates for differential evolution and PINNs are provided in Table 4, with the associated performance metrics (R^2^, MSE) summarized in Table 5. Both differential evolution and PINN come to similar optimizations for parameter values in most cases for both treatment categories.

**Table 4.**
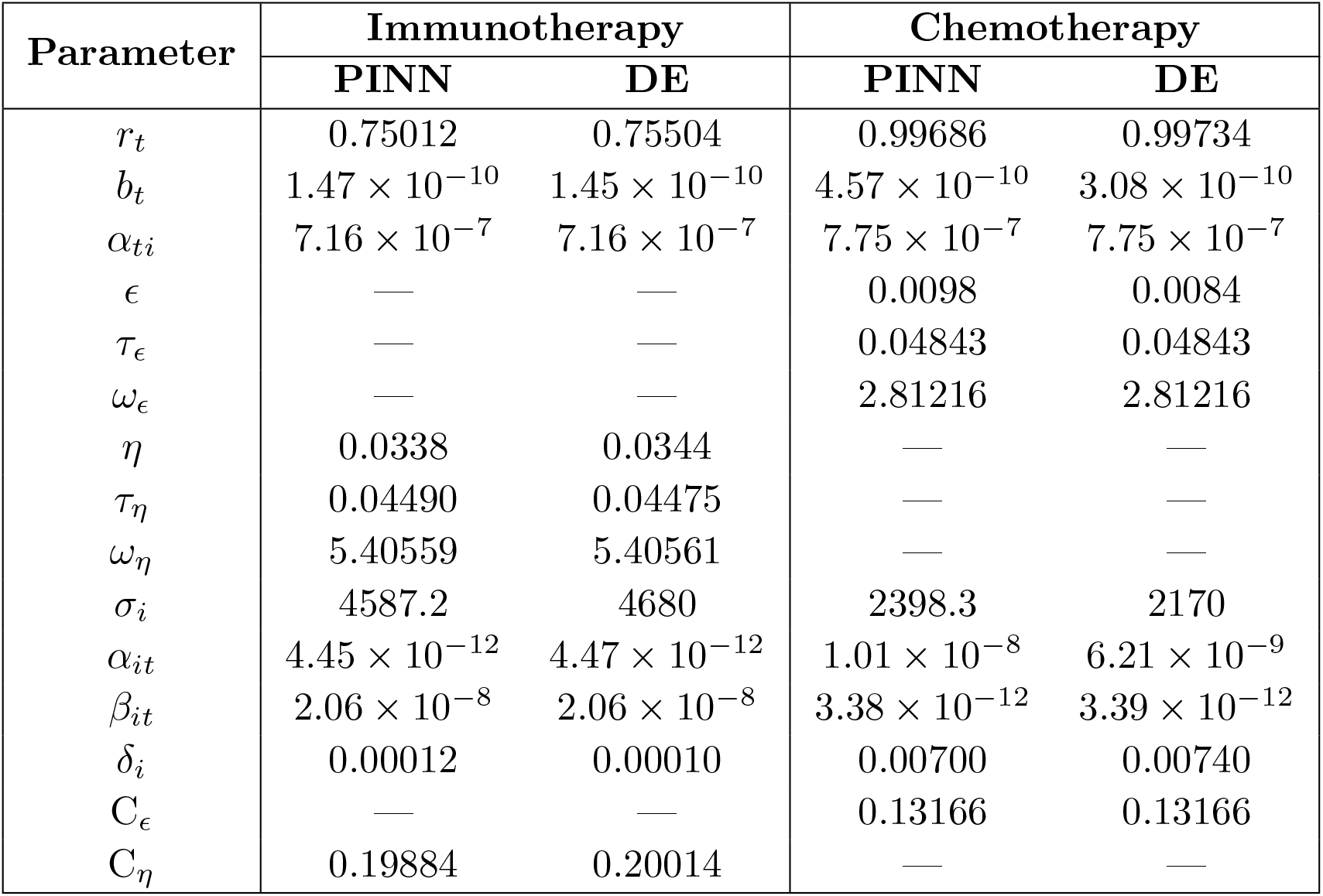
Comparison of estimated parameters used in Eqs. (3) – (8) from PINN and Differential Evolution for patients who received a curable outcome undergoing treatment with immunotherapy or chemotherapy.

**Table 5.** Comparison of R^2^ and MSE values for PINN and DE methods on immunotherapy and chemotherapy datasets for patients that experienced remission with therapy.

| Metric | Immunotherapy |  |  | Chemotherapy |  |  |
| --- | --- | --- | --- | --- | --- | --- |
| | PINN $\hat{T}(t; \theta^*)$ | PINN $\theta^* \rightarrow T(t)$ | DE | PINN $\hat{T}(t; \theta^*)$ | PINN $\theta^* \rightarrow T(t)$ | DE |
| R<sup>2</sup> | $9.9921 \times 10^{-1}$ | $9.9998 \times 10^{-1}$ | $9.9997 \times 10^{-1}$ | $9.9980 \times 10^{-1}$ | $9.9997 \times 10^{-1}$ | $9.9997 \times 10^{-1}$ |
| MSE | $4.6573 \times 10^{-5}$ | $1.0899 \times 10^{-6}$ | $1.5546 \times 10^{-6}$ | $1.1750 \times 10^{-5}$ | $1.7385 \times 10^{-6}$ | $1.7509 \times 10^{-6}$ |

**Fig 2.**
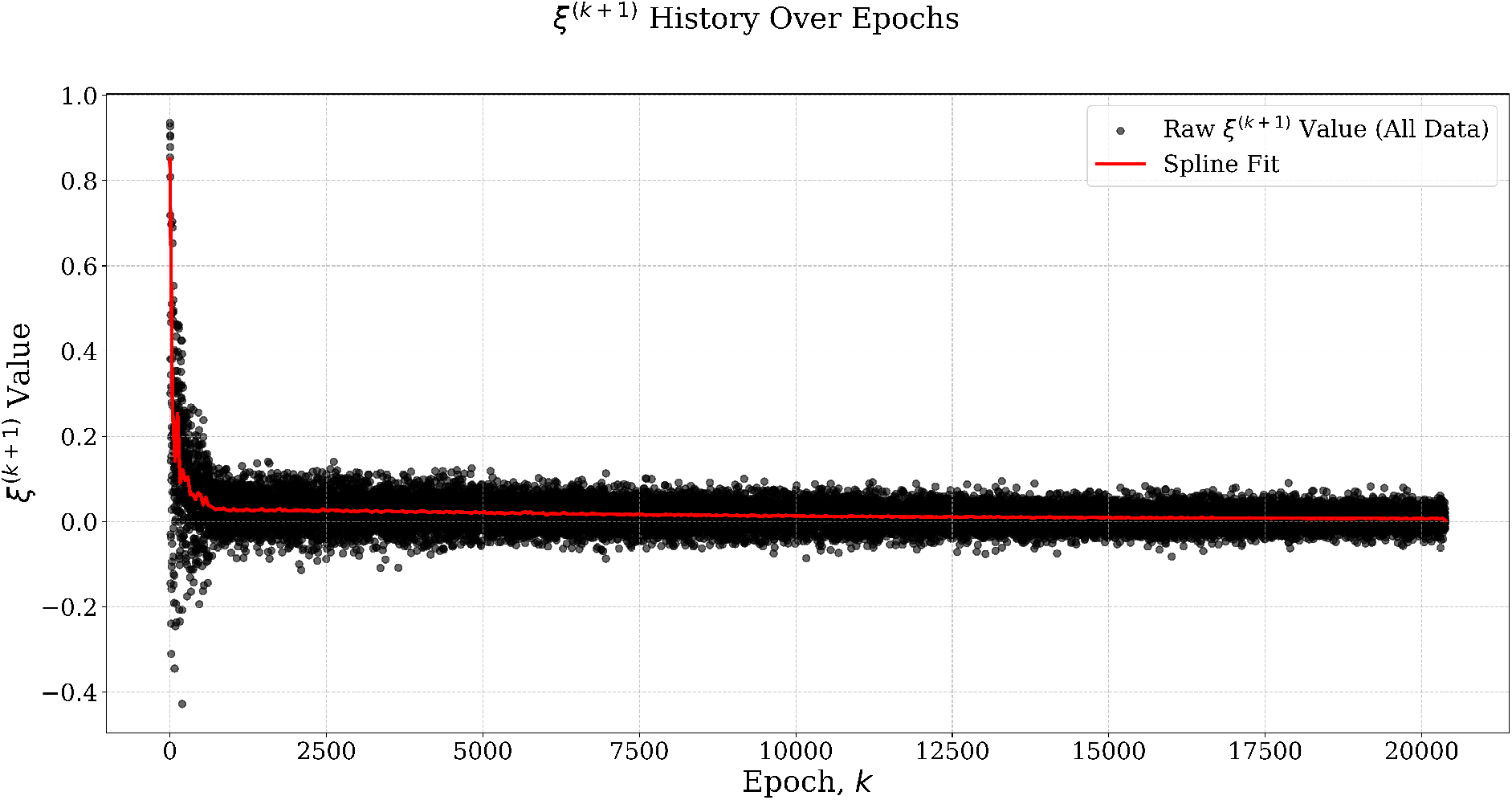
*ξ*^(*k*+1)^ stopping condition equation raw values plotted alongside a spline interpolation of the data downsampled the iterations done during training shows high divergence in the tumor estimation between epochs that reaches stability as the number of epochs increases.

Patients treated with immunotherapy who experienced remission of their cancer and their corresponding differential evolution and PINN models can be seen in Fig. 3a. The differential evolution fits the synthesized data very closely with an R^2^ value of 9.9997 × 10^−1^ and has an MSE of 1.5546 × 10^−6^. PINN improves on these parameter values and when the PINN parameter estimations are used in the SciPy ODE solver, a higher R^2^ value of 9.9998 × 10^−1^ and a lower MSE of 1.0899 × 10^−6^ are achieved. However, PINN’s model for solving the ODE and encoding the parameters on the model has slightly worse performance as the R^2^ value is lower at 9.9921 × 10^−1^ and the MSE is higher at 4.6573 × 10^−5^. The 95% pointwise confidence intervals were extremely narrow (average NBW = 2.4%, maximum NBW = 22.9%), resulting in low empirical coverage of 29.4%. This indicates that while the model provides highly precise predictions, it underestimates predictive uncertainty in this dataset.

**Fig 3.**
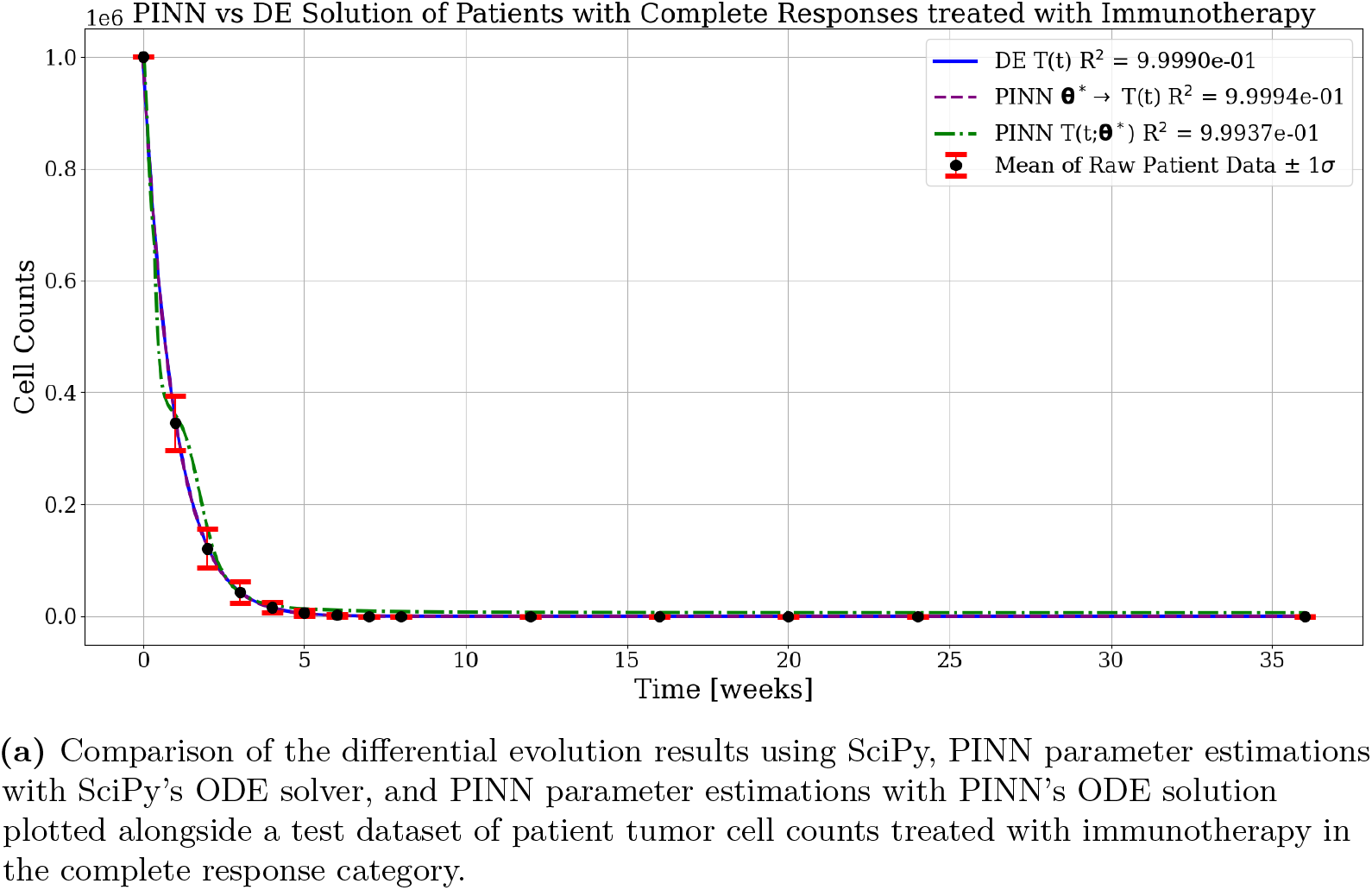
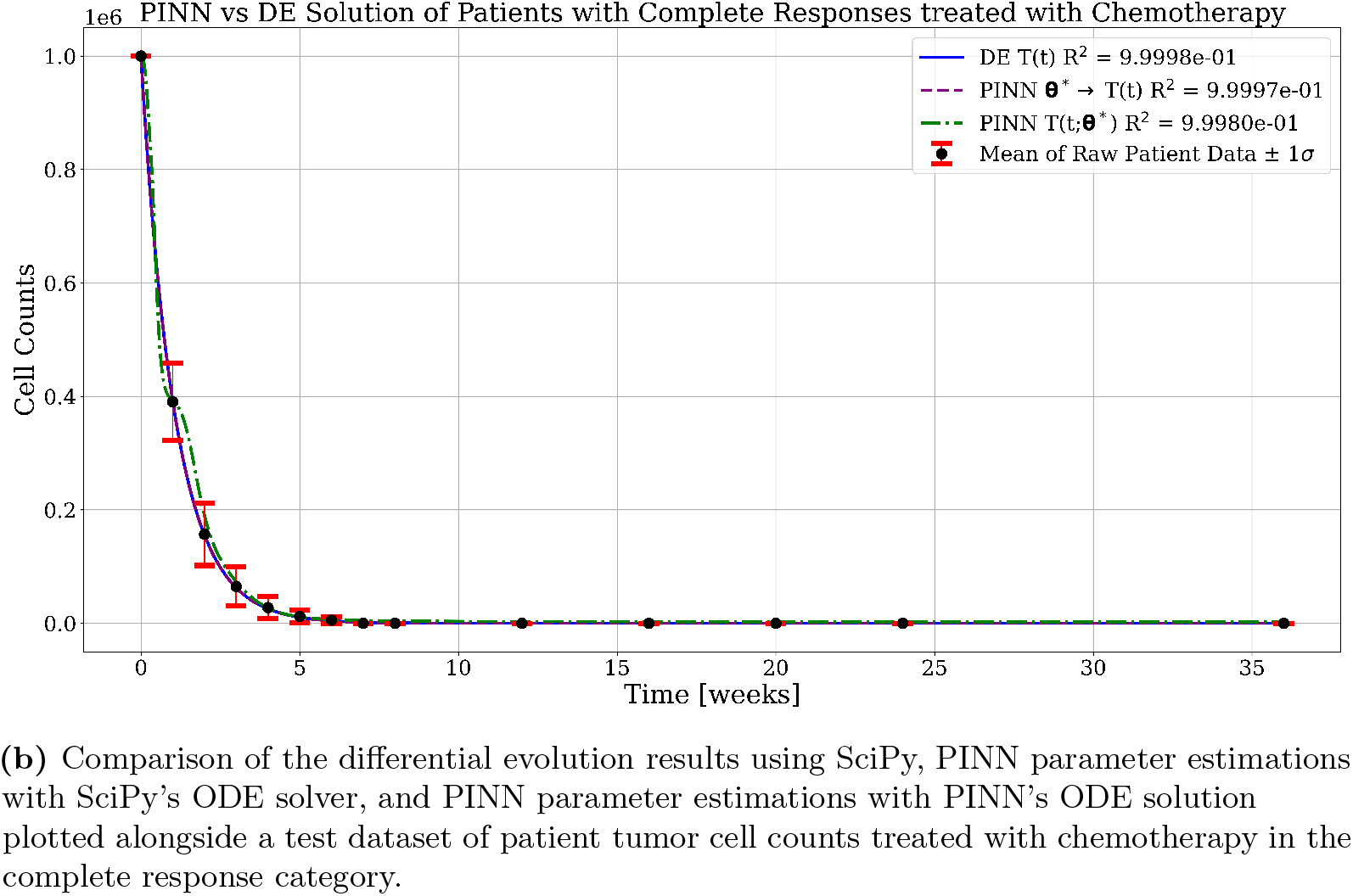
Comparison of differential evolution and PINN parameter and model estimations for tumor cell counts in complete response categories for immunotherapy and chemotherapy treatments.

Patients treated with chemotherapy who experienced remission of their cancer and their corresponding differential evolution and PINN models can be seen in Fig. 3b. The differential evolution fits the synthesized data very closely with an R^2^ value of 9.9998 × 10^−1^ and has an MSE of 9.6845 × 10^−7^. PINN iterates on these parameter values and when the new estimations are used in the SciPy ODE solver, an equal R^2^ value of 9.9998 × 10^−1^ and a statistically similar MSE of 9.6834 × 10^−7^ are achieved. However, like in the immunotherapy treatment cohort, PINN’s model for solving the ODE has slightly worse performance as the R^2^ value is lower at 9.9980 × 10^−1^ and the MSE is higher at 1.1750 × 10^−5^. Similar to the immunotherapy cohort, the 95% pointwise confidence intervals for the chemotherapy cohort were extremely narrow (average NBW = 1.2%, maximum NBW = 27.2%), resulting in low empirical coverage of 41.2%. This indicates that while the model provides highly precise predictions, it underestimates predictive uncertainty in this dataset.

In both treatment cohorts under the complete response scenario, the PINN achieves parameter estimates that are comparable to, and in some cases better than, those obtained with differential evolution on the synthetic datasets (Fig. 3). Nonetheless, when evaluating the model trajectories, the PINN’s neural solution exhibits slightly reduced accuracy relative to direct numerical integration with the SciPy ODE solver. This degradation is modest—the PINN still yields high R^2^ values—but becomes apparent when comparing relative R^2^ MSE across methods, where the SciPy solver consistently achieves lower error.

The modest discrepancy between the PINN and the direct SciPy ODE solution arises from the different approximation strategies employed. Whereas SciPy integrates the governing equations directly with adaptive numerical solvers that minimize local truncation error, the PINN learns an approximate mapping 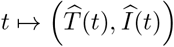 through a neural network. This introduces additional approximation error due to finite network capacity, optimization dynamics, and the competing balance between data-fitting and physics-based residual losses. As a result, even if the parameter estimates are accurate, the neural solution may not reproduce trajectories with the same precision as direct numerical integration, particularly in low-variance or noise-free regimes where solver accuracy is very high.

### Partial Response Cohort

The estimation of the parameters from differential evolution and PINN for patients who experienced a partial response for both types of treatments are listed in Table 6, subdivided into treatment categories, with corresponding performance metrics presented in Table 7. Both differential evolution and PINN made similar estimations for parameter values in most cases for both treatment categories.

**Table 6.** Comparison of estimated parameters used in Eqs. (3) – (8) from PINN and Differential Evolution for patients who had a partial response undergoing treatment with immunotherapy or chemotherapy.

| Parameter | Immunotherapy |  | Chemotherapy |  |
| --- | --- | --- | --- | --- |
|  | PINN | DE | PINN | DE |
| $r_t$ | 0.55968 | 0.57247 | 0.56375 | 0.54967 |
| $b_t$ | $8.70 \times 10^{-7}$ | $8.63 \times 10^{-7}$ | $9.29 \times 10^{-7}$ | $9.33 \times 10^{-7}$ |
| $\alpha_{ti}$ | $1.61 \times 10^{-7}$ | $1.61 \times 10^{-7}$ | $1.47 \times 10^{-7}$ | $1.49 \times 10^{-7}$ |
| $\epsilon$ | — | — | 0.04204 | 0.04350 |
| $\tau_\epsilon$ | — | — | 0.02926 | 0.02918 |
| $\omega_\epsilon$ | — | — | 8.71435 | 8.71435 |
| $\eta$ | 0.01229 | 0.0129 | — | — |
| $\tau_\eta$ | 0.02976 | 0.02974 | — | — |
| $\omega_\eta$ | 3.51577 | 3.51577 | — | — |
| $\sigma_i$ | 1995.7 | 2050 | 3631.0 | 3610 |
| $\alpha_{it}$ | $2.11 \times 10^{-8}$ | $2.46 \times 10^{-8}$ | $8.05 \times 10^{-12}$ | $8.05 \times 10^{-12}$ |
| $\beta_{it}$ | $1.32 \times 10^{-8}$ | $1.12 \times 10^{-8}$ | $1.49 \times 10^{-9}$ | $1.49 \times 10^{-9}$ |
| $\delta_i$ | 0.00093 | 0.00090 | 0.00159 | 0.00160 |
| $C_\epsilon$ | — | — | 0.10320 | 0.17512 |
| $C_\eta$ | 0.04381 | 0.10428 | — | — |

**Table 7.** Comparison of R^2^ and MSE values for PINN and DE methods on immunotherapy and chemotherapy datasets for patients that experienced a partial response with therapy.

| Metric | Immunotherapy |  |  | Chemotherapy |  |  |
| --- | --- | --- | --- | --- | --- | --- |
| | PINN $\hat{T}(t; \theta^*)$ | PINN $\theta^* \rightarrow T(t)$ | DE | PINN $\hat{T}(t; \theta^*)$ | PINN $\theta^* \rightarrow T(t)$ | DE |
| R<sup>2</sup> | $9.9833 \times 10^{-1}$ | $9.5241 \times 10^{-1}$ | $9.4322 \times 10^{-1}$ | $9.9898 \times 10^{-1}$ | $9.0697 \times 10^{-1}$ | $9.4488 \times 10^{-1}$ |
| MSE | $6.4848 \times 10^{-5}$ | $1.8444 \times 10^{-3}$ | $2.2005 \times 10^{-3}$ | $3.9478 \times 10^{-5}$ | $3.5911 \times 10^{-3}$ | $2.1277 \times 10^{-3}$ |

Tumor cell counts for patients treated with immunotherapy that experienced partial remission of their cancer are plotted along with differential evolution and PINN models in Fig. 4a. The differential evolution results in a relatively high R^2^ value of 9.5195 × 10^−1^ and an MSE of 1.8624 × 10^−3^. The parameter estimations from differential evolution are used as a starting point for PINN, which then encodes the parameters on its model as it tries to fit the data by meeting the constraints of the hyperparameters *λ*_data_, *λ*_phys_, and *λ*_init_ using an annealing scheduling for *λ*_phys_ defined in Eq. (42). Using the PINN’s parameter estimation with the SciPy’s ODE solver results in a higher R^2^ value of 9.5307 × 10^−1^ and a lower MSE of 1.8188 × 10^−3^. While this appears to be a modest improvement compared to the differential evolution estimate, when the full model output is used to estimate the data, the R^2^ value increases to a relative maximum of 9.9833 × 10^−1^ and the MSE reduces to a relative minimum of 6.4848 × 10^−5^. The mean trajectory *µ*(*t*) had an average normalized band width of 9.4% (max 23.5%), and the empirical coverage of observed test points within these intervals was 100.0%, indicating that the model provides well-calibrated local uncertainty estimates.

**Fig 4.**
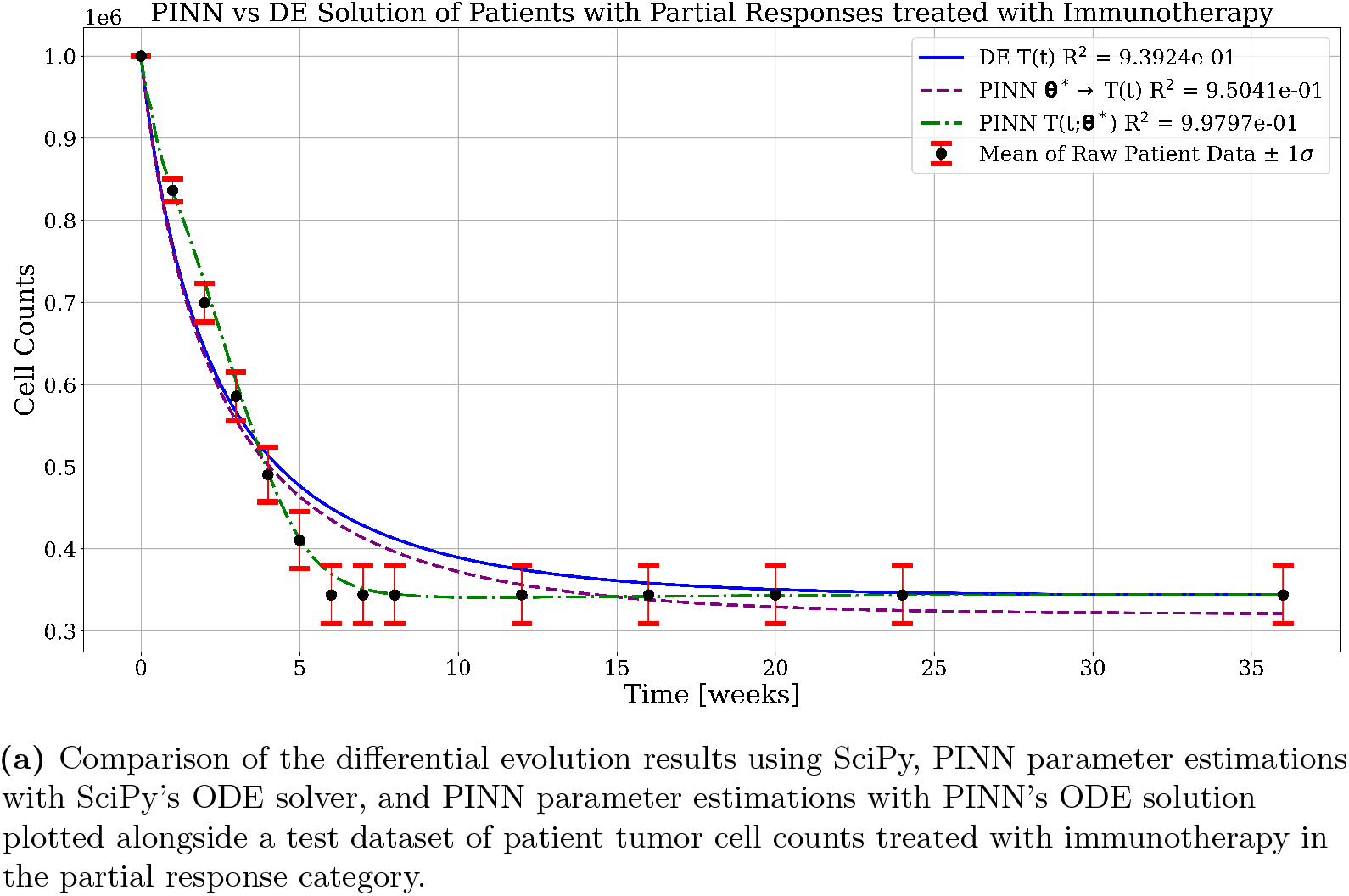
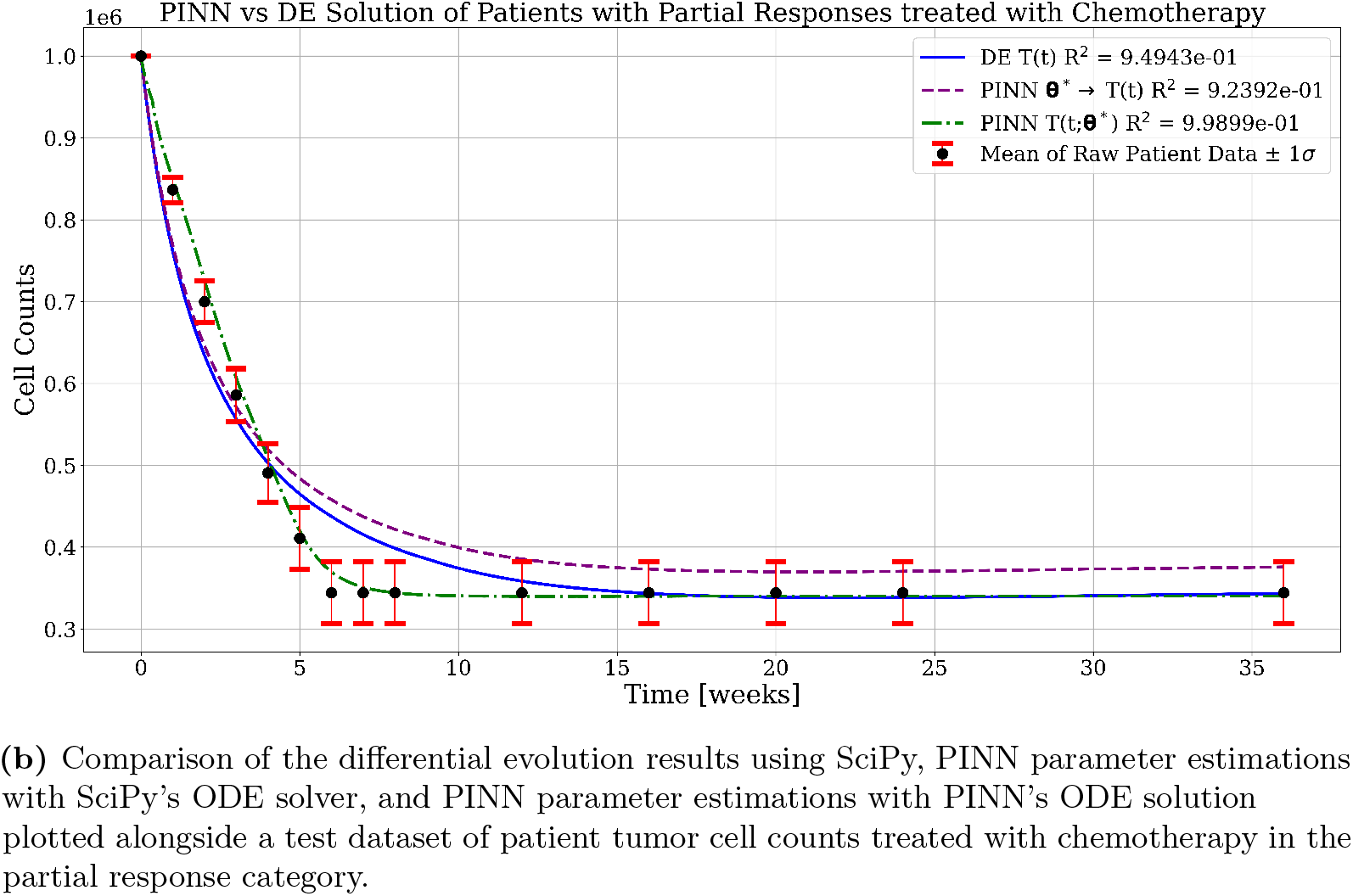
Comparison of differential evolution and PINN parameter and model estimations for tumor cell counts in partial response categories for immunotherapy and chemotherapy treatments.

The chemotherapy-treated cohort that experienced partial remission can be seen in Fig. 4b, grouped together with the differential evolution and PINN models that fit the data. The differential evolution results in an R^2^ value of 9.5259 × 10^−1^ with an MSE of 1.8301 × 10^−3^. Using the output of the PINN estimation of the parameter values with the SciPy ODE numerical solver results in a slightly diminished R^2^ value of 9.4968 *×* 10^−1^ and a higher MSE of 1.9426 *×* 10^−3^. The PINN model has the greatest fit of the data, with the highest R^2^ value of 9.9898 *×* 10^−1^ and the lowest MSE of 3.9478 × 10^−5^. The mean trajectory *µ*(*t*) had an average normalized band width of 7.1% (max 17.9%), and the empirical coverage of observed test points within these intervals was 100.0%, indicating that the model provides well-calibrated local uncertainty estimates.

A consistent pattern across cohorts is that parameter estimates produced by the PINN do not always translate to improved performance when evaluated solely with a conventional numerical integrator. This discrepancy arises from the differing objectives: differential evolution directly minimizes a data-only loss using the numerical integrator, whereas the PINN jointly optimizes network weights and parameters under a composite loss that balances data fidelity, ODE residuals, and initialization constraints. The PINN’s flexible function approximator can partially absorb model–data mismatch, thereby allowing parameter values that reduce the combined PINN loss but are not the minimizers of the integrator-only MSE. Consequently, in some cohorts the PINN-derived parameters produce superior integrator-based fits, while in others they do not — a result that reflects the multi-objective nature of PINN training rather than a failure of parameter estimation per se. We therefore caution against treating a single-objective numerical solver (e.g., SciPy’s **solve_ivp**) as a full cross-validation of PINN parameters unless the PINN training explicitly enforces equivalence between the network solution and the numerical integrator (for example, by embedding the integrator within the training loop or by placing stronger weight on the physics term), which was not the case for our training loop.

## Conclusion

In this work, we proposed a dynamical model of tumor–immune interactions with therapeutic input terms that capture the build-up and decay of drug action through a convolutional dosing scheme, closely mimicking clinical treatment schedules.

To address the scarcity of cellular-resolution clinical data, we introduced a Monte Carlo-based algorithm to synthesize realistic longitudinal datasets conditioned on patient outcomes. This approach opens new possibilities for generating data-driven models in oncology, where direct cellular measurements are limited.

We then applied differential evolution (DE) for parameter estimation, providing a strong baseline for data-driven model fitting. These estimates were used as initialization for a physics-informed neural network (PINN), which jointly optimized parameters and trajectories under a composite loss function balancing data fidelity, ODE residuals, and initialization constraints. To improve training stability, we adapted a decorrelation metric from maximum-likelihood estimation and proposed it as a novel stopping criterion, showing that training could be halted once residual correlations diminished across epochs.

Our results demonstrate that PINNs can serve as a powerful alternative to traditional optimization-based parameter estimation. In low-variance datasets, PINN-derived parameters produced lower MSE and higher R^2^ than DE when evaluated with a single-objective solver, though the neural solution itself exhibited modest degradation. In higher-variance datasets, PINN parameters had mixed performance under solver evaluation, but the full PINN model consistently outperformed both baselines, achieving substantial reductions in MSE and increases in R^2^. These findings underscore a key distinction: while DE identifies parameters that minimize solver-based error, the PINN optimizes a multi-objective loss in which the network itself can absorb part of the model-data mismatch.

When analyzing the confidence intervals of the Monte Carlo-generated trajectories, we observed that low-variance trajectories often yielded low empirical coverage, whereas high-variance trajectories consistently achieved perfect empirical coverage.

This discrepancy is likely due to the use of a single, fixed dropout rate across datasets. Achieving both well-calibrated empirical coverage for the Monte Carlo trajectories and accurate coverage of the mean data points may require tuning the dropout rate on a per-dataset basis during training and evaluation.

In summary, this work bridges mathematical modeling, machine learning, and cancer research by (i) proposing a realistic therapeutic model, (ii) introducing a novel data synthesis pipeline, (iii) adapting a decorrelation-based stopping criterion for PINN training, and (iv) benchmarking DE and PINN methods across treatment cohorts. The approach highlights both the promise and limitations of PINNs for biomedical parameter estimation: they offer strong data fits and flexibility, but their parameter values are not always directly transferable to single-objective numerical solvers. Future research should focus on integrating single-objective solver-based objectives into PINN training and applying the method to other clinical datasets to evaluate its translational potential.

Specifically, to address the computational demands and “parameter transferability” issues noted in this study, ongoing work extends the methodologies developed here into two distinct optimization frameworks. First, we are incorporating the decorrelation-based early stopping criterion as a trigger for gradient-informed polishing within a novel GPU-accelerated Differential Evolution algorithm. This hybrid approach aims to bridge the gap between global search and local refinement, using the PINN’s stability metrics to decide when to transition from exploration to gradient-based exploitation.

Furthermore, the structural complexity of these tumor–immune models suggests a need for optimization strategies that can handle both nonconvexity and specific structural constraints. To this end, we are currently expanding our optimization suite through the development of a GPU-accelerated Population-Based Great Deluge (PBGD) algorithm. By applying PBGD to both black-box and structure-aware optimization problems, we aim to further enhance the scalability and robustness of parameter estimation in complex biological systems, leveraging high-performance computing to achieve high-fidelity fits across even larger, more heterogeneous patient cohorts.

## Acknowledgments

The views and opinions expressed in this article are those of the authors and do not necessarily reflect the official views or positions of PeopleTec. The authors thank the PeopleTec Technical Fellows for providing a review of our paper prior to submission.

